# The push-pull intercrop *Desmodium* does not repel, but intercepts and kills pests

**DOI:** 10.1101/2022.03.08.482778

**Authors:** Anna Laura Erdei, Aneth Bella David, Eleni C. Savvidou, Vaida Džemedžionaitė, Advaith Chakravarthy, Béla Péter Molnár, Teun Dekker

**Affiliations:** Department of Plant Protection Biology, Swedish University of Agricultural Sciences, Alnarp, Sweden; Department of Zoology, Plant Protection Institute, Centre for Agricultural Research, ELRN, Budapest, Hungary; Department of Molecular Biology and Biotechnology, University of Dar-es-Salaam (UDSM), Dar-es-Salaam, Tanzania; Department of Agriculture Crop Production and Rural Environment, University of Thessaly, Volos Greece

## Abstract

Over two decades ago, scientists developed a push-pull intercropping strategy that received critical acclaim for synergizing food security with ecosystem resilience in smallholder farming. The strategy suppresses Lepidopteran pests in maize through a combination of a repellent intercrop (push), commonly *Desmodium* spp., and an attractive, dead-end border crop (pull). Key is the intercrop’s constitutive release of volatiles that repel herbivores. Surprisingly, however, we found that *Desmodium* does not constitutively release volatiles, and only minimally upon herbivory. Further, in oviposition choice settings, *Spodoptera frugiperda*, a devastating invasive pest, was not repelled by *Desmodium* volatiles. In search of an alternative mechanism, we found that neonate larvae strongly preferred *Desmodium* over maize. However, their development stagnated and none survived. In addition, larvae were frequently seen impaled and immobilized by the dense network of silica-fortified, non-glandular trichomes. Thus, entirely different from repelling adult moths, *Desmodium* intercepts and decimates dispersing offspring. As a hallmark of sustainable pest control, maize-*Desmodium* intercropping has inspired countless efforts trying to emulate a stimulo-deterrent diversion in other cropping systems. However, detailed knowledge of the actual mechanisms is required to rationally improve the strategy, and translate the concept into other cropping systems.

## Main text

Since the dawn of agriculture, humanity has been in an arms race with insect pests. Traditionally, a set of integrated cultivation strategies tailored to local settings helped keeping pests at bay, including associational resistance through varietal mixtures and intercropping^1–3^. With the advent of agrochemicals, monocultures superseded traditional strategies. However, their profound externalities on ecosystem resilience and global climate^4,5^ have resuscitated interest in more sustainable alternatives, frequently grafted on traditional strategies. Trending terms such as agroecology, and climate smart, regenerative or organic agriculture evidence the search for solutions that harmonize food production and pest control with ecological sustainability. Some innovative practices have been important sources of inspiration. Among these, the push-pull strategy in which maize is intercropped with the legume, *Desmodium*, is arguably the most well known^6^.

Push-pull aims to reduce the abundance of insect pests in crops through repelling the pest in the crop, while simultaneously providing attractive sources to trap the pest out (formalized by Miller and Cowles^7^). Using this ‘stimulo-deterrent diversion’ principle, a push-pull strategy was devised to combat Lepidopteran pests in sub-Saharan smallholder maize farming^8,9^. Embroidering on the common practice of smallholder farmers to intercrop maize with e.g. edible pulses, the strategy uses the perennial fodder legume *Desmodium* as intercrop in maize plots. *Desmodium* reportedly constitutively releases large amounts of terpenes (such as (*E*)-4,8-dimethyl-1,3,7-nonatriene ((*E*)-DMNT), (*E*)-*β*-ocimene and cedrene) that repel (‘push’) lepidopteran pests and attract natural enemies (‘pull’)^10–12^. A ‘dead-end’ host sown as border crop (another ‘pull’ component), typically napier grass, complements the strategy as it induces oviposition in Lepidoptera, but reduces larval survival compared to maize^11–13^. This cropping strategy reduces infestations of various Lepidoptera pests, including *Chilo partellus* and *Busseola fusca*, as well as *Spodoptera frugiperda*, a polyphagous invasive pest that is ravaging maize and vegetable production and threatens food security in sub-Saharan Africa^14,15^. Strongly propagated by institutions and governments^16–21^, this intercropping strategy has found widespread adoption in East Africa. As a hallmark of sustainable pest control, it also serves as a tremendous source of inspiration for intervention strategies in other cropping systems.

The ‘push’ volatiles reported in previous studies^11,12^ are typically released by plants after induction by herbivory. This begs the question of why *Desmodium* releases these volatiles constitutively. Push-pull maize-*Desmodium* intercropping causes substantial shifts in below-ground ecosystems, including increased soil microbe diversification, increased soil nitrogen and carbon, increased plant defense through plant-soil feedback, and suppression of parasitic weeds and pathogenic microbes^22,23^. We therefore verified if the ‘constitutive’ release of volatiles was, in fact, induced or enhanced by soil-borne interactions. The root-microbe interactions are of particular interest, given the intimate association of legumes with specific microbial groups e.g. rhizobia and mycorrhizae. Indeed, soil and root-microbe interactions can induce pathways that lead to release of volatiles^e.g., 22,24^

Surprisingly, however, *D. intortum*, which is by far the most commonly used intercrop in push-pull technology^10^, did not release volatiles constitutively at all (Figure 1a, b, Extended Data, Figure 2 and 3). This was independent of the soil in which *D. intortum* was grown, whether live soil (organic potting soil, organic clay Swedish soil or African clay loam soil from *D. intortum* plots), autoclaved soil, or autoclaved soils inoculated with mycorrhiza or rhizobacteria (Extended Data, Figure 4, 5 and 6). None of the previously reported terpenes^12^ were constitutively released, nor any terpene or other volatiles that are typically released upon herbivory. Similar results were obtained with *D. uncinatum* (Extended Data, Figure 7). In contrast, we did confirm that *Melinis minutiflora*, a Poaceae used previously as a push intercrop, constitutively releases a diverse blend of terpenes in large quantities (Extended Data, Figure 2, 3 and 8). Clearly, independent of soil interactions, *Desmodium* does not constitutively release volatiles.

**Fig. 1:**
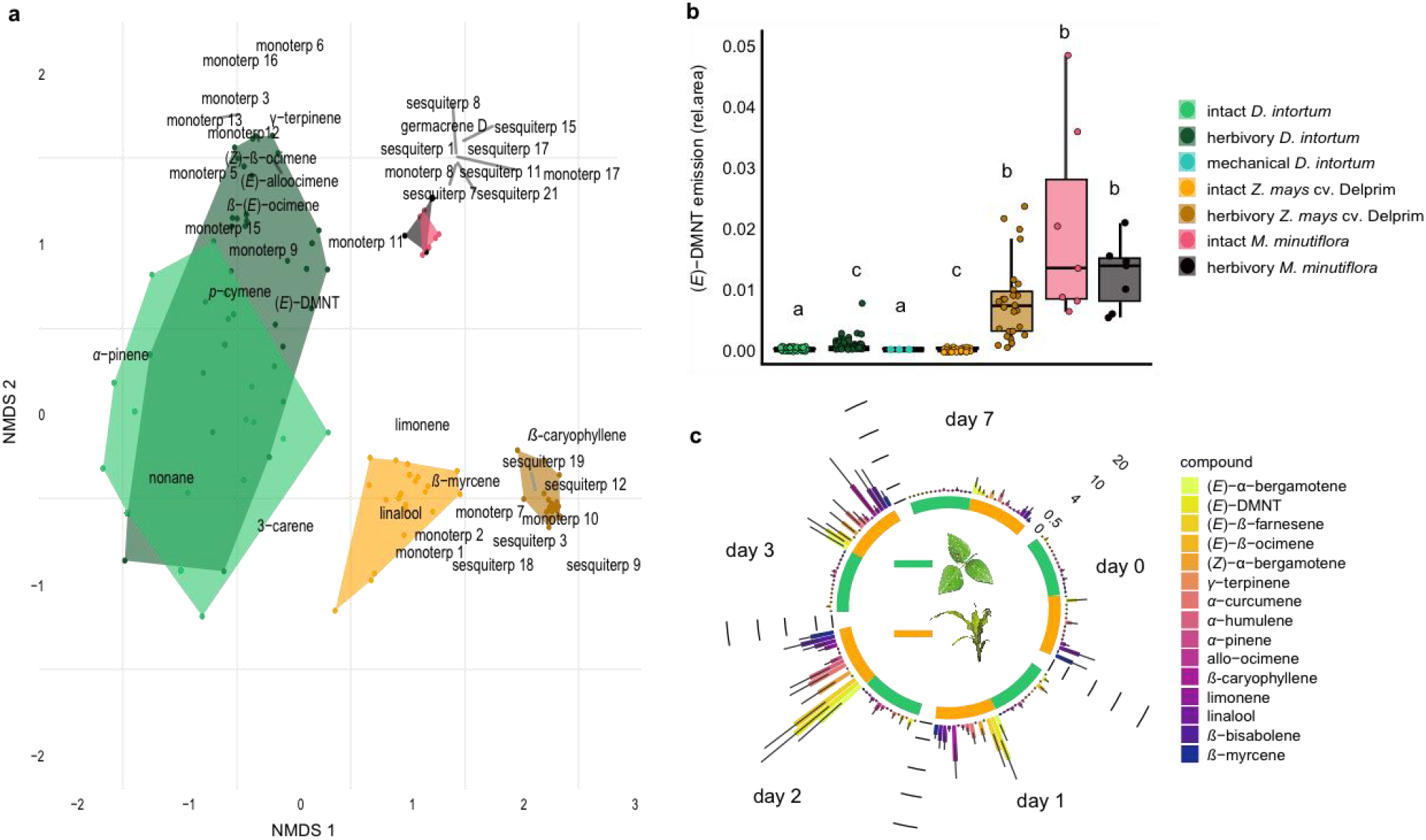
*Desmodium intortum* does not constitutively release terpene volatiles, and hardly following larval feeding. **a**, Nonmetric multidimensional scaling (NMDS) analysis of volatiles emitted by *D. intortum, Z. mays* cv. Delprim and *M. minutiflora* plants, intact and 48 hrs following *S. frugiperda* feeding (stress value = 0.138). (*E*)-4,8-dimethyl-1,3,7-nonatriene ((*E*)-DMNT), (*Z*)-*β*-ocimene, (*E*)-*β*- ocimene and (*E*)-alloocimene were not constitutively released, and only in low quantities in response to herbivory. Volatiles emitted by intact and herbivore-induced *D. intortum* (F_model_ = 15.597, R^2^ = 0.132, *p*_adj_ = 0.021) and *Z. mays* plants (F_model_ = 50.521, R^2^ = 0.512, *p*_adj_ = 0.021) were significantly different in PERMANOVA and pairwise comparison, but emissions from intact and herbivore induced *M. minutiflora* plants (F_model_ = 1.469, R^2^ = 0.109, *p*_adj_ = 1) were not. **b**, (*E*)-DMNT emission before and 48 hrs following herbivory (n = 8, ± SE). The absolute peak areas were divided by the peak area of the internal standard and divided by the sum of monoterpenoids across all laboratory volatile collections for normalization. Treatments with different letters are different (Kruskal-Wallis with Benjamini and Hochberg *p* value correction, χ^2^ = 57.315, *p* =1.578 10^-10^). **c**, Emission of volatile monoterpenoids and sesquiterpenoids from *D. intortum* and *Z. mays* before, during and after *S. frugiperda* larval feeding (n = 5, ± SE). Peak areas of each terpenoid were divided by the area of the internal standard and divided by the sum of monoterpenoids or sesquiterpenoids across all laboratory volatile collections. Error-bars show the standard error for relative volatile emission of each group. Day 0 - volatile emission before herbivory, Day 1 - 24 hrs after herbivory, Day 2 after 48 hrs, and so on. Larvae were removed after 48 hrs.

Although the constitutive release of volatiles is an important precondition for push-pull, inadvertent herbivory of *Desmodium* could have induced volatile release reported in earlier studies. However, *D. intortum* only minimally released induced volatiles when either mechanically damaged or when fed upon by *S. frugiperda* larvae (Figure 1a-d, Extended Data, Figure 2 and 3). This contrasted with maize, which, in line with previous studies^25–27^, released large amounts of herbivore-induced volatiles in response to herbivory, with emission peaking between 24 and 48 hrs following infestation, and declining over the course of 7 days (Figure 1c). Herbivory of *M. minutiflora* did not significantly boost release of volatiles above the already high constitutive release (Figure 1b, Extended Data, Figure 2 and 3).

Arguably, greenhouse conditions are not representative of field conditions and additional, unknown factors in the field may cause the release of volatiles by *Desmodium*. We therefore analyzed 50 headspace samples from *D. intortum* from seven locations in Tanzania and Uganda. Also under field conditions, terpene release by *D. intortum* was minimal (Figure 2, Extended Data, Figure 8), and possibly induced by herbivory that was visible on most sampled plants. Thus, regardless of whether constitutive or induced, *Desmodium* does not release terpene volatiles, or any other volatiles, in large quantities in the field. Although it cannot be excluded that other conditions or herbivores may induce higher release of reported volatiles, our data with numerous samples under different growth conditions, and from different geographic regions show that this must be very rare, and can therefore not be at the core of a generic strategy. In contrast, maize, all of which displayed some herbivore damage, did release typical herbivore induced volatiles^25,26^ (Fig 2, Extended Data, Figure 8), with variations likely due to differing levels of and age since herbivore infestations, which could not be controlled in the field.

**Fig. 2:**
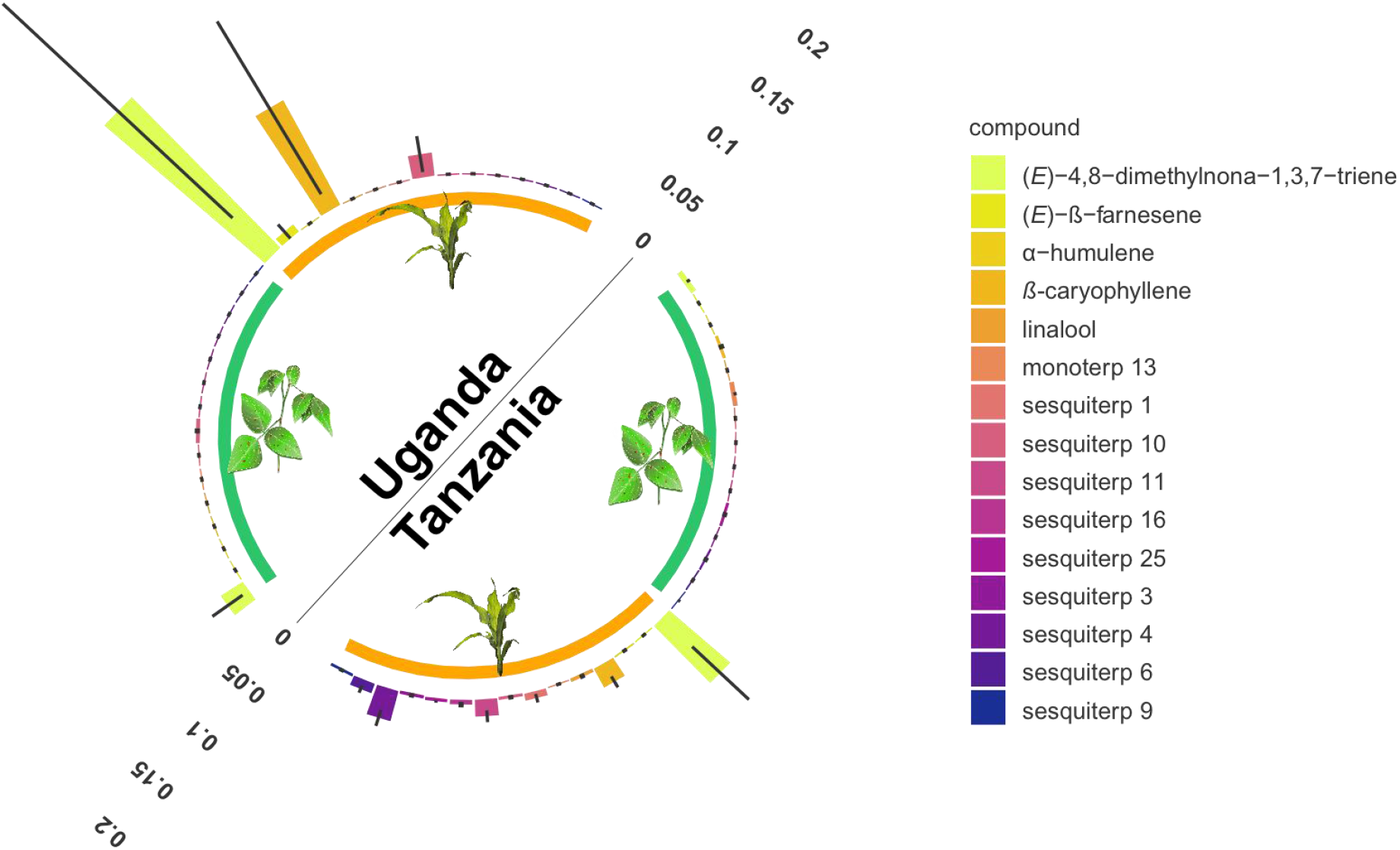
Monoterpenoid and sesquiterpenoid emission by *D. intortum* and *Zea mays* plants under field conditions at several locations in Tanzania and Uganda. The absolute peak area of each peak was divided by the sum of the area of monoterpenoids or sesquiterpenoid emission across all samples from the same location. Error bars represent ± SE on the scale of the relative volatile emission. Minor terpenoid compounds were not identified to species level as this was not the focus of the study, and was further hindered by the vast diversity of compounds and the lack of synthetic standards.

Ironically, if the mode of action in maize-*Desmodium* push pull was repellent terpene volatiles, induced maize itself would appear a much better push candidate than *Desmodium*. Although the lack of volatiles emitted made it highly unlikely that *Desmodium* repels lepidopteran pests, we double checked this in bioassays. In a wind tunnel, gravid *S. frugiperda* were given a choice between maize plants with either *D. intortum* or artificial plants in the background (Extended Data, Figure 1). Adult females landed and oviposited on either maize plant equally, underlining that *D. intortum* volatiles indeed did not repel gravid *S. frugiperda* (Figure 3c).

**Fig. 3:**
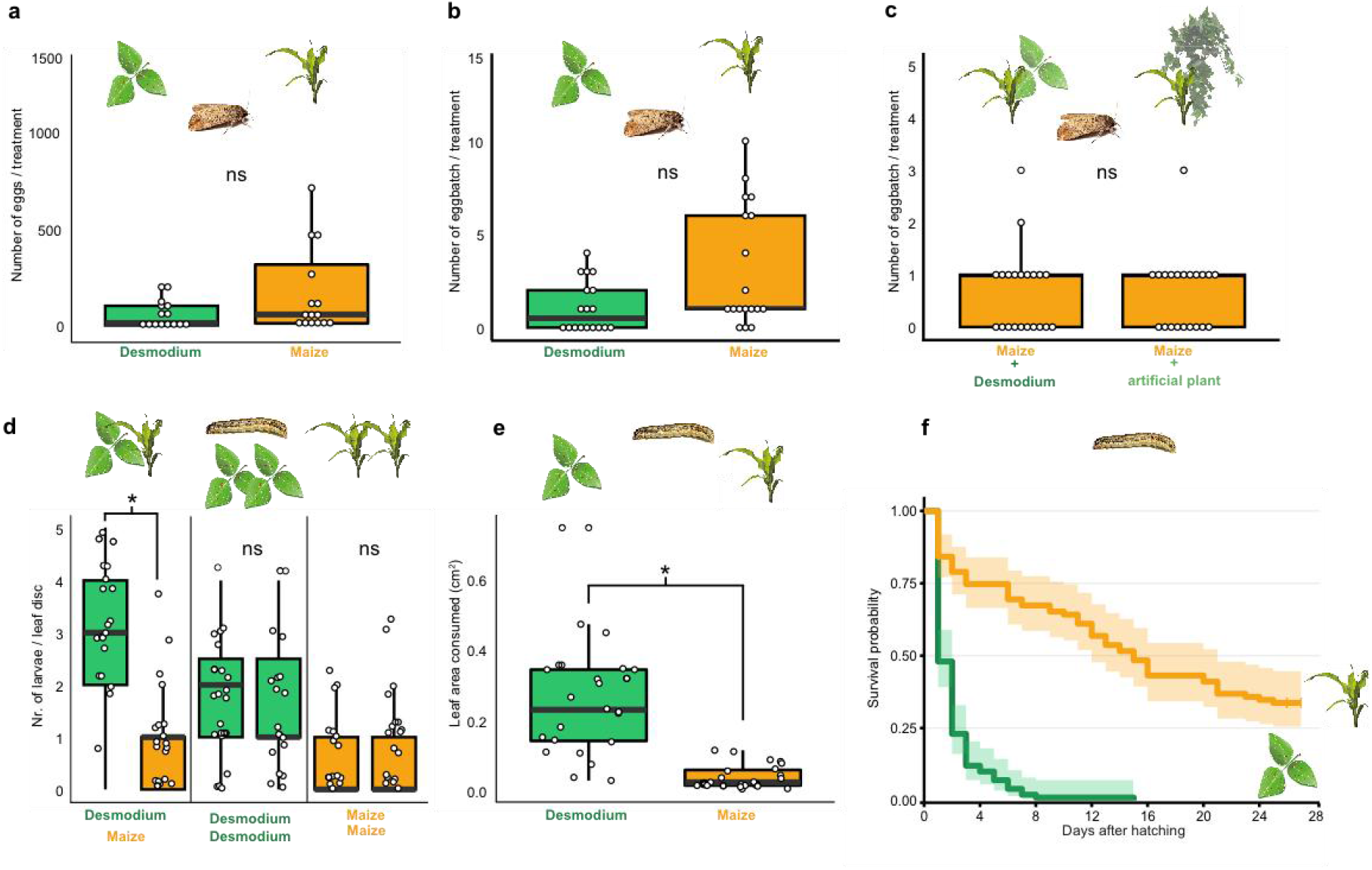
*D. intortum* does not repel ovipositing *S. frugiperda*. Instead it is prefered by larvae but truncates their development. **a**, The number of eggs laid on *D. intortum* or *Z. mays* plants in choice-experiments in cages (n = 25) did not differ (Wilcoxon signed rank exact test, *p* = 0.055). **b**, Number of egg batches laid on *D. intortum* or *Z. mays* plants (n = 25, Wilcoxon signed rank exact test, *p* = 0.075). **c**, Number of egg batches on *Z. mays* plants in a background of either *D. intortum* plant or a plastic plant mimic did not differ in wind tunnel oviposition assays (n = 21, Wilcoxon signed rank exact test, *p* = 0.825). **d**, First instar *S. frugiperda* larvae preferred *D. intortum* against *Z. mays* in two choice leaf disc bioassays (n = 25, Wilcoxon signed rank exact test, *p* = 2.73*10^-3^). **e**, First instar *S. frugiperda* larvae consumed more *D. intortum* than *Z. mays* (20 hrs, two-choice leaf disc bioassays, n = 25, Wilcoxon signed rank exact test, *p* = 3.338*10^-6^). **f**, Survival probability of *S. frugiperda* on diets consisting of *D. intortum* (greenleaf *Desmodium*) was lower than on *Z. mays*, with no larvae surviving on *D. intortum*. (Kaplan-Meier survival analysis, *p* = 2.000*10^-16^). Error bars, ± SE.

Evidently, to explain the suppression of lepidopteran pests using *Desmodium* as intercrop, one needs to invoke a different mechanism than ‘stimulo-deterrent diversion’ or ‘push-pull’. To investigate possible alternatives we scored female *S. frugiperda* oviposition preference, larval feeding preference, and larval survival on maize and *Desmodium*. First, in two-choice tests *S. frugiperda* preferred oviposition on maize over *Desmodium*. However, the preference was not strong, as females also oviposited on *Desmodium*. In the field, one could perhaps expect a further shift toward *Desmodium*, particularly when maize is small and *Desmodium*, a perennial, well developed. However, irrespective of female oviposition choice, many lepidopteran larvae are known to disperse from the plant on which they hatched. Neonate larvae typically ‘parachute’ between plants using silk threads^28–30^, whereas later larval stages actively disperse across the soil surface in search for new host plants^30–32^. Given the dense, continuous ground cover of *Desmodium* in the interrows, stochastically the large majority of dispersing larvae would end up in *Desmodium*, particularly when maize plants are small and *Desmodium*, a perennial, large. We therefore verified the preference and survival of *S. frugiperda* larvae on *Desmodium* compared to maize. Surprisingly, first instar larvae strongly preferred *D. intortum* over maize, both in choice and in leaf area consumed (Figure 3d,e). However, their development stagnated, with hardly any larva molting to the second instar, and none completing their development (Figure 3f, Extended Data, Figure 9).

In addition to stagnating development, we found that larvae, particularly later larval instars, moved slowly on *Desmodium* leaves and stems, while many were immobilized entirely. Closer scrutiny of *D. intortum* surfaces revealed a dense network of non-glandular, uniseriate and uncinate trichomes, with densities and a distribution depending on the surface type (Figure 4a - d, f, Extended Data, Figure 10a). The stems and main veins of the leaves were particularly densely populated with uncinate trichomes. First instar larvae were somewhat freely moving and grazing between trichomes (Extended Data, Figure 10b,c), but older larvae were seen impaled and immobilized by these trichomes (Figure 4c,d, Extended Data, Figure 10d-f). Occasionally, even ovipositing *S. frugiperda* were immobilized with their ovipositor on *D. intortum* (Extended Data, Figure 10g). Whereas trichomes were flexible at the base, they were fortified with silica toward the tip (Figure 4f), equipping the plant with an effective mechanism to obstruct, damage and immobilize herbivores. Also beneficial insects (Extended Data Figure 10i) and even vertebrates can be trapped by *Desmodium^33^*. Similar structures are also used by many other plant species^34–36^, and may serve multiple purposes including seed dispersal^37,38^.

**Fig. 4:**
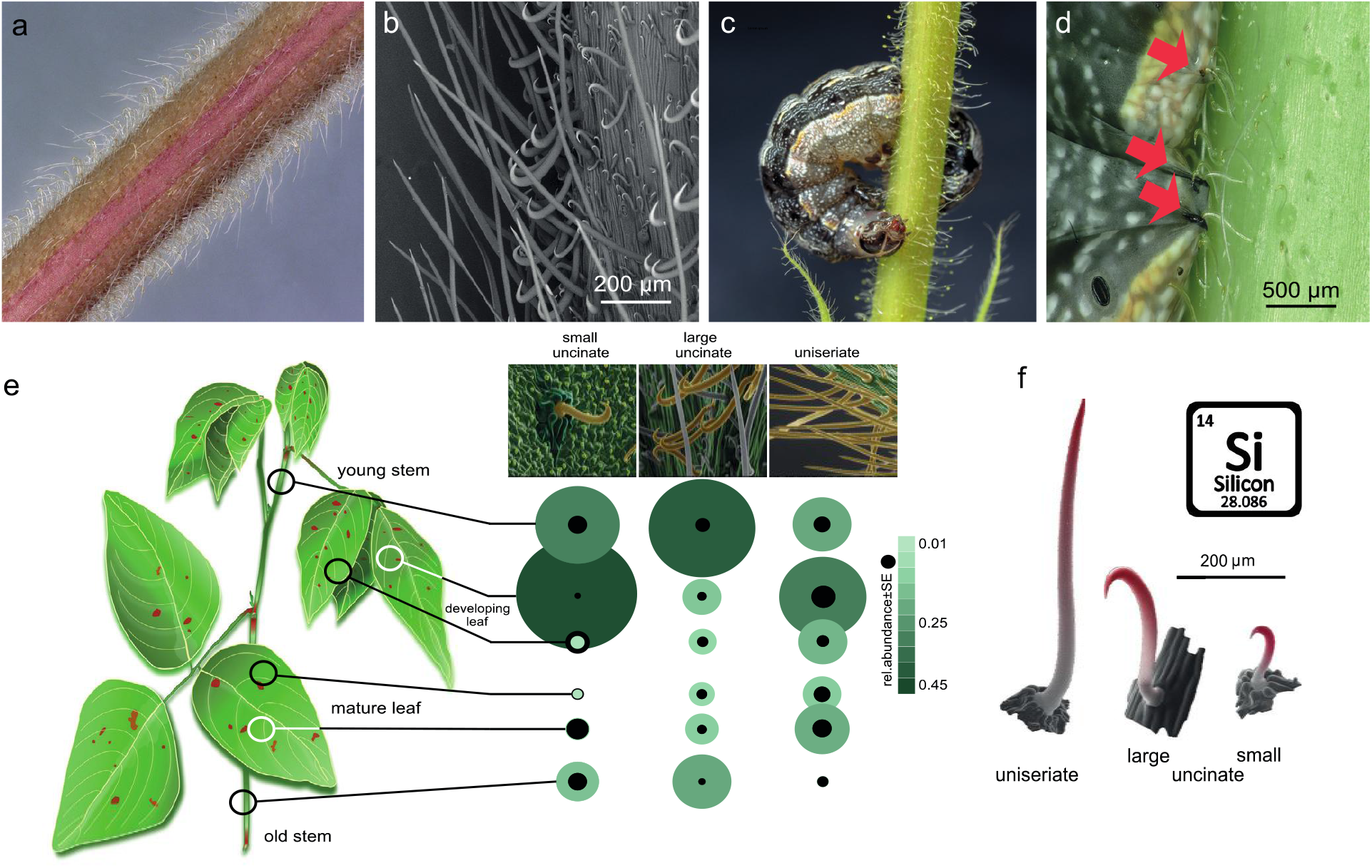
Non-glandular trichomes on *Desmodium intortum* act as a physical barrier for herbivores. **a**, Light microscopy image of a section of a young *D. intortum* stem densely covered with trichomes. **b**, Scanning electron microscopy (SEM) image of a young *D. intortum* stem. Straight uniseriate hairs (up to 2 mm long) extended beyond the large (0.2 - 0.4 mm) and small (0.05 - 0.2 mm) hooked uncinate trichomes (scale bar: 200 μm). **c,** A fifth instar *S. frugiperda* larva impaled and immobilized on a stem of *D. intortum* by both large and small uncinate trichomes. **d**, Fourth instar *S. frugiperda* larva pierced by uncinate trichomes (red arrows). Trichomes either immobilized larvae or broke off from the basal cell with the tip remaining in the larval body causing severe wounds. **e**, Distribution of non-glandular trichomes on different parts of the *D. intortum* plant. The relative abundance was calculated as the mean of trichome count divided by the sum of trichomes per trichome type across samples. Black circles indicate the standard error of relative trichome abundance (n = 5). **f**, SEM images combining EDX element topography images indicate relative surface silica (Si) distribution (red) of uniseriate, large and small uncinate trichomes (n = 5).

We thus infer that in the field *Desmodium* affect fitness of lepidopteran larvae, both directly and indirectly. First, *Desmodium* entices larval feeding, but truncates larval development. Second, trichomes on *Desmodium* hinder movement, damage the cuticle and even entirely immobilize larvae on the plant, increasing developmental time, exposure to natural enemies and overall mortality^39,40^. Third, the ingestion of trichomes will damage the intestinal lining and affect digestion, development and survival^40,41^. Indeed, while first instar larvae easily fed around the trichomes, larger larvae did ingest trichomes as evidenced by trichomes found in larval frass. Effectively, rather than functioning as a repellent intercrop, *Desmodium* appears to be a developmental deathtrap for larvae.

Clearly ‘push’ does not describe the mode-of-action of *Desmodium*. Instead, the plant exhibits properties reminiscent of a ‘pull’ crop, a ‘dead-end host’. Although superficially similar in mode of action to the ‘pull’ border crop Napier grass, *Desmodium* is distinctly different, as it is preferred by larvae, not by adults^8,10^. In addition, *Desmodium* forms a mechanical barrier to dispersing larvae. Further field studies need to detail how oviposition preference, larval dispersal, development and survival on *Desmodium*, mechanical obstruction by *Desmodium*, and additional mechanisms such as parasitization and predation, interplays with crop phenology in suppressing various lepidopteran species across the cropping season. Knowing the exact interaction of mechanisms is critical if we for instance wish to substitute the fodder crop *Desmodium* with a food crop to enhance food security, or if we are to translate the concept of interceptive intercropping to other cropping systems.

The surprising discovery that *Desmodium* hardly emits volatiles and does not repel herbivores contrasts strongly with the very large number of publications and the huge global attention that maize-*Desmodium* push-pull technology has garnered over more than two decades. Indeed, the narrative of the ‘push’ crop *Desmodium* repelling moths has been mentioned by numerous papers since its first mention around the year 2000. Astonishingly, however, close scrutiny of the literature revealed a total absence of primary data. Whereas the most cited paper from around 2000, Khan and colleagues^12^, mentions some of the *Desmodium* volatiles and claims repellence of stemborers, no primary chemical analytical or behavioral data were presented in this paper, nor in any preceding or ensuing paper. Equally remarkable is how, in spite of thousands of citations and an abundance of efforts to emulate push-pull in other cropping systems, this crucial detail has collectively slipped the attention of the scientific community.

Further research should study how pest suppression in interceptive intercropping is affected by factors such as pest species, natural enemies, crop phenology, insect population dynamics, and abiotic factors including soil and climate, and others. This will be pivotal for improving the current maize intercropping strategy, tailoring it to the needs of local smallholder farmers and other ecosystem services sought after (e.g. replacing *Desmodium* with food crops with similar properties^34–36,41–43^), as well as rationally translating the concept to other cropping systems.

## METHODS

### Plants

Seeds of the most common intercrop species in push-pull farming (*Desmodium intortum*, greenleaf *Desmodium*, and *Desmodium uncinatum*, silverleaf desmodium) were acquired from Simlaw seeds Co. Ltd, Nairobi, Kenya). *M. minutiflora* seeds were obtained from the South African Sugarcane Research Institute (SASRI, Mount Edgecombe, South Africa). Maize seeds (*Zea mays* cv. Delprim) were provided by the laboratory of Ted Turlings at University of Neuchâtel, Switzerland. The cultivar is a European commercial hybrid and long-time standard whose volatile emission patterns have been thoroughly studied^44^.

*Desmodium* spp. seeds were sterilized by using 3% NaOCl and rinsed in distilled water and germinated on wet filter paper, and transferred to seedling trays with live or autoclaved soil (121 °C for 20 min). After 21 days the plants were transferred to 18 cm diameter pots containing live or autoclaved soil and were grown for 8 weeks in a greenhouse (22 – 25 °C, light cycle 16:8 hrs, RH 65%). Another set of plants were raised from cuttings of mature stem parts of *D. intortum* and rooted in distilled water. Rooted cuttings were then planted in pots containing autoclaved soil with different inoculants: 200 g soil of a Tanzanian push-pull field per each pot, autoclaved soil with 60 mg of *Rhizobium leguminosarum, Bradyrhizobium japonicum* mixture per each pot (equal portions of *Rhizobia* inoculant for *Phaseolus* beans, and soy beans from Samenfest GmbH., Freiburg, Germany) or autoclaved soil with 120 mg of mycorrhizal fungi inoculate per each pot (mixture of *Glomus intraradices*, *G. etunicatum*, *G. monosporum*, *G. deserticola*, *G. clarum*, *Paraglomus brasilianum*, *Gigaspora margarita*, *Rhizopogon villosulus*, *R. lutcolus*, *R. amylopogon*, *R. fulvigleba*, *Pisolithus tinctorius*, *Scleroderma cepa* and *S. citrinum*, Wildroot Organic Inc., Texas). The microbial inoculants were premixed in autoclaved soil before plant inoculation. Plants from cuttings grown on autoclaved soil were used as control. *M. minutiflora* seeds were germinated in live soil in plastic trays, and the seedlings were transferred into pots with live soil after two sets of leaves appeared. Eight weeks old *M. minutiflora* and *Desmodium spp*. plants were used in the experiments. Maize seeds were planted directly into live or autoclaved soil in pots and maintained in the greenhouse for 6 weeks.

For the cage oviposition experiments, maize seeds were sown next to 5 weeks old *D. intortum* plants in 12 cm pots and grown together for three weeks. For the wind tunnel experiments, maize and *D. intortum* plants were grown in separate pots and four to five weeks old maize and nine to eleven weeks old *D. intortum* plants were used.

### Insect rearing

*S. frugiperda* were obtained from the Ted Turlings laboratory at University of Neuchâtel, Switzerland, and were raised on a soybean based semi artificial diet supplemented maize whorls. The third instar larvae were separated into groups of ten individuals in plastic boxes.

Pupae were sexed and separated in rearing cages. Adults were provided with a 5 % sucrose solution and 6 days old adults were mated for 6 hrs and used in oviposition experiments.

### Volatile collections

The plants grown in the greenhouse were enclosed in a 60 cm x 20 cm polyethylene (PET) oven bag (Toppits ^®^ ‘Bratschlauch’, Melitta, Minden, Germany) above ground for 24 hrs to saturate the headspace. Prior to sampling, 2 μl of 250 ng/ul nonane solution in hexane was injected onto a piece of filter paper into the oven bag 40 minutes prior to sampling. Solid phase microextraction (SPME) fibers (DVB/CAR/PDMS 50/30 μm, Supelco, Sigma-Aldrich, Bellefonte, PA, USA) were conditioned at 250 °C in the split/splitless injector of the GC-MS in split mode for 10 minutes. The SPME fibers were exposed to the closed headspace for 30 minutes. The volatile emission of intact, mechanically damaged and herbivore-damaged plants were sampled. *D. intortum* plants were mechanically damaged by cutting ten randomly selected leaflets in half, perpendicularly to the midrib. For herbivore-treatment, eight fourth to fifth instar and 12 hrs starved *S. frugiperda* larvae were put on the plants. In the first sets of experiments the feeding period lasted for 48 hrs before volatile sampling.

A time series experiment of volatile terpenoid emission following herbivory was performed on *D. intortum* and *Z. mays* cv. Delprim plants grown on autoclaved soil inoculated with Tanzanian soil. Eight fourth instar larvae were put on each plant after 12 hrs of starving and removed after 48 hrs of feeding. The plants were sampled before herbivory and after 24 hrs, 48 hrs of herbivory. Larvae were removed from the plants after 48 hrs and plants were resampled 72 hrs and one week after the start of the experiment. The volatile headspace was closed for 24 hrs before each sampling and the SPME sampling procedure was the same as described above.

Field volatile samples of *D. intortum* (greenleaf *Desmodium*) and *Z. mays* were collected on farmer fields in Tarime and Musoma districts in Mara region, Tanzania, and Rural Community in Development (RUCID) center, in Mityana district, Uganda. Healthy *D. intortum* plants and maize plants with visible herbivore damage were selected and enclosed in 60 cm x 20 cm polyethylene (PET) oven bags for 18 hrs overnight. The use of standard and the SPME volatile sampling procedure was the same as described above.

### Gas chromatography coupled mass spectrometry (GC-MS)

A GC-MS (Agilent technologies, 7890B GC coupled with 5975 MSD) was used for SPME analysis. Fibers were inserted into a 250 °C splitless injection port with The split valve closed for 1 min. The GC was equipped with a DB-WAX column (60 m x 250 μm x 0.25 μm). The carrier gas was helium and the total column flow was 34.883 mL/min. The oven temperature was programmed as follows: 50 °C/min, 10°C/min to 220 °C, 20 °C/min to 250 °C. The final temperature was held for 1 min. The mass spectrometer was used in electron ionization mode 70 eV and the detector scanned in the 29-400 m/z range. Samples were also injected on a GC-MS equipped with an HP-5 column (Agilent technologies, 6890 GC coupled with 5977A MSD, column: 60 m x 250 μm x 0.25 μm), with similar inlet settings and carrier gas (helium). The oven program was as follows: 40 °C/2 min, 8 °C/min to 230 °C. The solvent delay and mass spectrometry settings were the same as described above.

GC-MS results were analyzed using Agilent Mass Hunter B.08.00, the peaks were auto integrated with agile integrator and manual integration. Compounds were tentatively identified by matching their mass spectra with those found in MS Libraries (NIST11 and Wiley12). The identification was verified by comparing calculated Kovats retention indices (RI) to those published in the NIST WebBook database and PubChem database and comparisons with analytical standards (See list of synthetic compounds in Table S1).

### Oviposition choice experiments

We conducted two experiments to study the short-range/multimodal oviposition repellency and long-range/olfactory oviposition repellency of *D. intortum* for *S. frugiperda* females.

#### Short-range/multimodal oviposition repellency experiments

In short-range/multimodal oviposition repellency experiments, maize seeds (*Z. mays* cv. Delprim) and *D. intortum* cuttings were co-planted. The experiments were conducted three weeks after co-planting, when the biomass of each plant were roughly similar. Plants were placed in 30 × 30 × 30 cm net cages (Bugdorm, Megaview, Taiwan) in a climate chamber set to 25±2 °C, 65%±5% relative humidity and 16:8 h L:D light cycle. Six days old virgin *S. frugiperda*, one female and one male, were mated for 6 hrs and females were let to oviposit for 48 hrs. A cotton ball soaked in 5% sucrose solution was placed between the plants for adult feeding. The egg batches and the number of eggs per each batch were counted at the end of the second day on both plants and the cage surfaces.

#### Long-range/olfactory oviposition repellency experiments

To score for spatial repellency of *D. intortum*, a modified wind tunnel (180 cm × 80 cm × 60 cm, 30 cm/s airflow) was used (Extended data, Figure 1). At the furthest upwind part of the flight section of the tunnel, two six-weeks old maize plants (*Z. mays* cv. Delprim) were positioned at 60 cm from each other. Directly upwind and separated by a stainless steel gauze (100 mesh) an eight-weeks old *D. intortum* or artificial plastic plant was placed directly upwind from the maize plants. In both sections a 20 cm plexiglass sheet was placed in line with the airflow to separate the airflow of the two sides (Extended data, Figure 1). Two six days old females and one six days old male were released in the chamber 1 hr prior to scotophase. A cotton ball soaked in 5% sucrose solution was placed in the chamber at the release side as a source of food. The position of the female and the number of egg batches laid on each side of the chamber were recorded after scotophase, 12 hrs following the start of the experiment.

### Larval choice experiments

We conducted two-choice feeding bioassays to determine the feeding preference of the first larval instar of *S. frugiperda*. We cut 8 mm diameter leaf discs from young leaves of 6-7 weeks old maize plants and leaves of 10-12 weeks old *D. intortum* plants. We put the leaf discs on wet filter paper discs 60 mm apart from each other in 100 mm x 20 mm plastic Petri-dishes. Ten one-day old *S. frugiperda* larvae were placed in each arena and the position of larvae was recorded after 1 h, 2 h and 20 h periods. After 20 h feeding each leaf disk was photographed and the consumed surface area of each disk was determined by image analysis using ImageJ (version 1.53)^45^.

### Larval survival experiments

Larval survival on maize and *D. intortum* scored in plastic petri-dishes (100 mm x 20 mm), which were lined with wet filter paper to increase humidity. Five first instar *S. frugiperda* larvae were moved to each arena on the day of egg-hatching and fed daily with an excess amount of freshly cut *D. intortum* leaves or leaf blades of 4-5 weeks old maize (*Z. mays* cv. Delprim). After reaching the fourth instar stage, the maize diet was supplemented with the ligule, leaf sheets and young stems of maize and the larvae were separated into individual plastic cups to prevent cannibalism. The growth of the larvae was monitored daily and we determined the larval stage based on body coloration and the diameter of head capsules. We terminated the experiment after the insects pupated.

### Light microscopy of *Desmodium* spp

Upper and mid stem branches as well as the leaves of healthy 8 weeks old *D. intortum* plants were sampled for light microscopy. In addition, *S. littoralis* larvae that were immobilized on *D. uncinatum* and *D. intortum* stems and leaves were observed and photographed with a digital light microscope (Keyence VHX-5000, Keyence Corporation, Osaka, Japan) equipped with standard zoom lens (VH-Z20R magnification: 20-200x and VH-Z100R magnification: 100-1000x). For detailed, high depth-of-field images, photo stacking technique was used. Series of images were captured (50-100 depending on the size of the examined larvae) at different focus distances (step size, 20 - 40 μm). Subsequently, partially focused images were combined with Helicon Focus software (Helicon Soft Ltd., Kharkiv, Ukraine) into a high depth of field image.

### Scanning electron microscopy of *Desmodium* spp

To get further insights in the structure of the *D. intortum* trichomes, scanning electron microscopy (SEM) was performed on leaf and stem samples. Healthy leaves and stems were collected from eight-weeks old and one-year old plants from the greenhouse, and scanned using a FEI Quanta 3D scanning electron microscope operating with a field emission gun (FEG) electron source, equipped with SE (LVSED/ETD), BSE (vCD) and EDAX SDD EDS detectors. Low vacuum mode (50-80 Pa specimen chamber pressure) was used in order to avoid sample charging, and allowed us to use plant material without sample fixation, dehydration and sample coating. The accelerating voltage was 10-20kV with 40-480 pA beam current.

Furthermore the elemental composition of trichomes was studied using energy-dispersive X-ray spectroscopy (EDX), acquisition time: 50 sec. Measurements were taken in four regions (base, lower and higher middle and tip) on the longer type of trichomes and from three regions in case of small uncinate trichomes.

### Statistical analysis

In case of each volatile sample the absolute peak areas were divided by the area of the internal standard peak to account for differences in volatile sampling efficiency. The volatile components were categorized into four compound groups: monoterpenoids, sesquiterpenoids, green leaf volatiles and other volatiles. We calculated the total sum of peak areas for these volatile groups across samples for the laboratory volatile collections and field volatile collections by location. The volatile collections were further normalized across samples by dividing the absolute peak areas by the sum of the total area of the volatile group from the corresponding dataset.

The clustered heatmaps of volatile emission profiles were generated from z-scores calculated from the normalized volatile data using package pheatmap^46^. Jaccard dissimilarity indices were calculated from binary (presence/absence) standardized volatile data and non-metric multidimensional scaling (NMDS) was completed using the metaMDS function of package vegan in R^47^. Permutational multivariate analysis of variance (PERMANOVA) was completed on Jaccard dissimilarity indices using the adonis function of the vegan package. For assessing differences in the normalized volatile peak areas for (*E*)-DMNT and (*E*)-*β*-ocimene between groups Kruskal-Wallis tests and Wilcoxon rank sum tests were used from package stats with Benjamini and Hochberg *p* value correction^48^.

We used Wilcoxon paired rank sum tests with a null hypothesis of random choice using package stats for two-choice oviposition experiments and larval choice experiments^48^. As the statistical power of Wilcoxon paired rank sum tests are limited, we also fitted generalized linear mixed models (GLMM) by maximum likelihood with fixed factor for choice and random factor for replication on the two-choice oviposition data using package lme4^49^. We used the simulation-based test from package DHARMa^50^ to assess the goodness of fit for the complete model. The post hoc tests were completed with the emmeans package using Tukey’s comparisons^51^.

Survival probabilities were calculated with Kaplan–Meier survival analysis^52^ and the survival curves were compared using a log-rank test between diets in package survival^53^. Survival curves were visualized using package survminer^54^.

## Data availability statement

Volatile analysis data associated with volatile analysis and behavioral bioassays are available in figshare with the identifier(s) [10.6084/m9.figshare.19297730] and GC-MS raw data from the authors upon reasonable request.

## ACKNOWLEDGEMENT

We are grateful to Mr. Samuel Nyanzi at Rural Community in Development (RUCID) center, Mityana, and Dr. Fred Kabi of Makerere University for support with volatile collections in Uganda. We thank Ms. Eva Svensson from Lund University for help with soil autoclave sterilization. We are thankful to Alex Berg for pictures of *Desmodium* spp. highlighting their trichomes as well as immobilized insect larvae and Ábel Szabó for his support in scanning electron microscopy. Special thanks to Prof. Ted Turlings’ lab for providing maize (*Zea mays*) seeds and *Spodoptera frugiperda* colonies and Prof. Peter Anderson for *Spodoptera littoralis* used in some pictures. We thank Prof. Marie Bengtsson, SLU, Sweden for providing standards for identification purposes. The following funding agencies are acknowledged for making this project possible: Sida, Food Security Program (ABD, TD), Ekhagastifelsen (ALE), SLU Global (TD), the EU Erasmus Program (ES), János Bolyai Research Scholarship of the Hungarian Academy of Sciences (BPM).

## AUTHOR CONTRIBUTIONS

ALE, ABD and TD conceived the idea and designed the experiments. All the authors contributed at different stages to performing the experiments, data analysis and writing of the manuscript.

## EXTENDED DATA

**Fig. 1:**
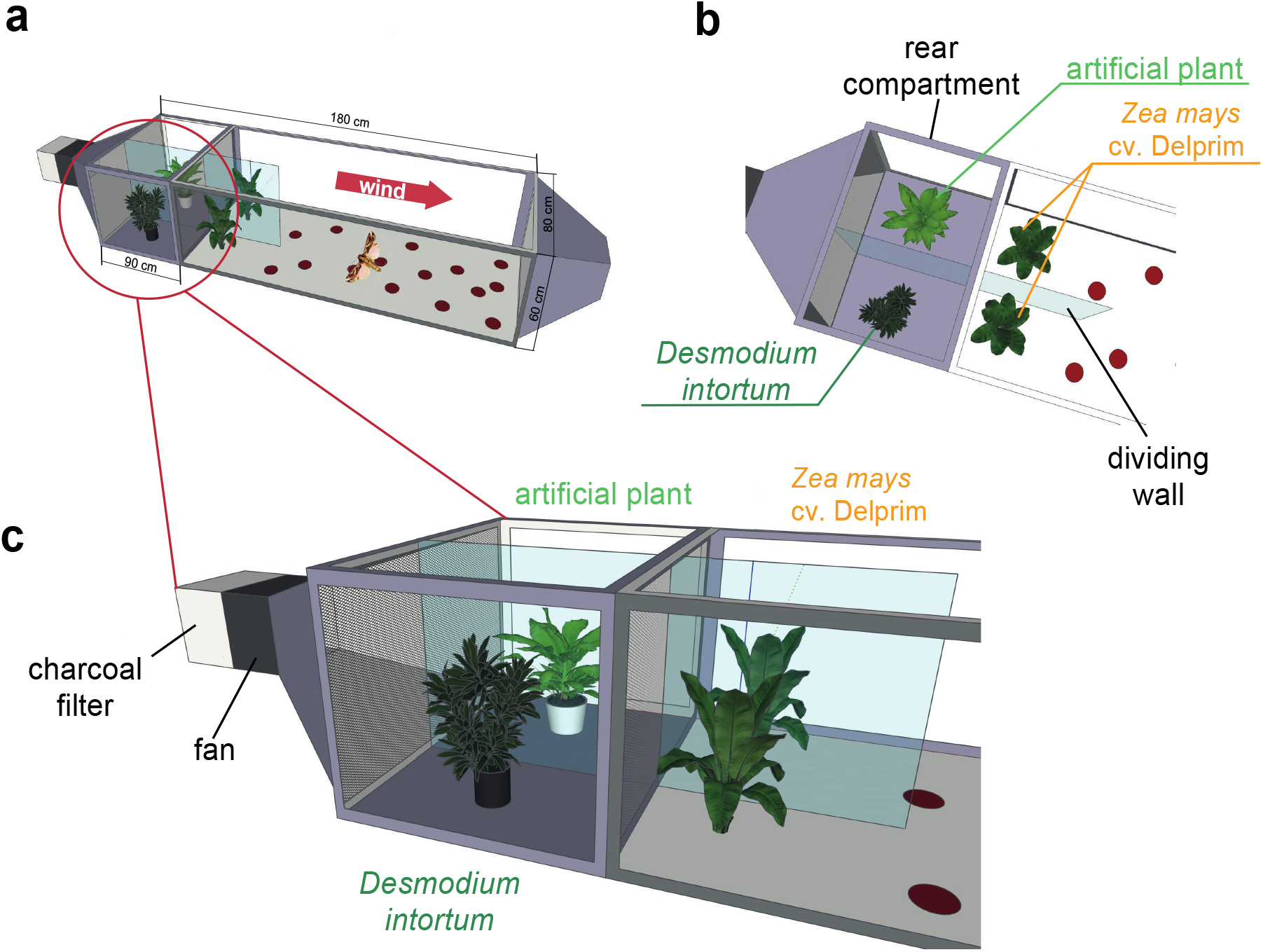
Wind tunnel setup to study the oviposition repellency of *Desmodium intortum* volatiles. Two *Zea mays* cv. Delprim plants were placed in laminar filtered air flow with *D. intortum* (greenleaf *Desmodium*) or a plastic mimic plant directly upwind from the flight chamber containing two maize plants. A gravid *Spodoptera frugiperda* female was released in the wind tunnel. The number of egg batches laid on both maize plants were counted and the position of mimic plants and *D. intortum* plants were randomized.

**Fig. 2:**
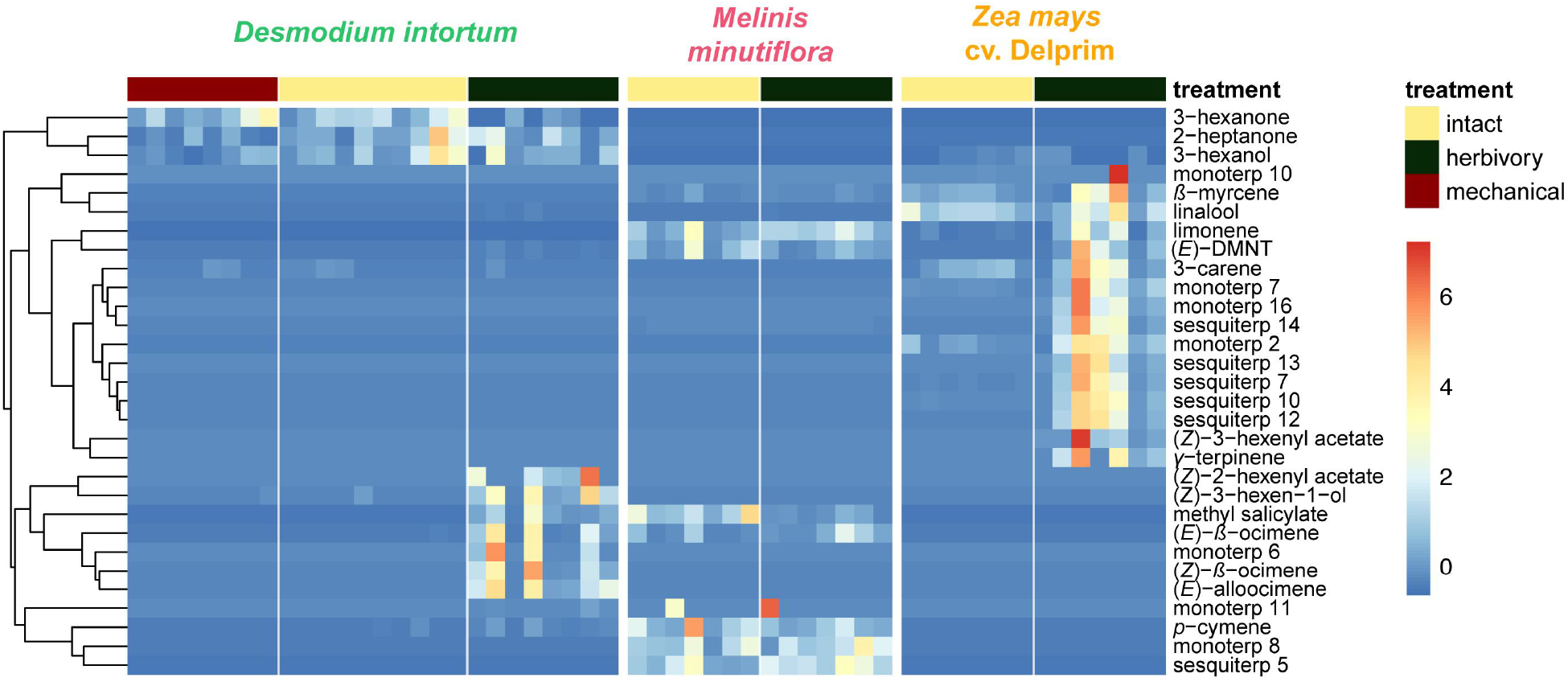
Heatmap showing relative amounts of headspace volatile compounds emitted from intact, herbivore induced and mechanically damaged *Desmodium intortum, Zea mays* cv. Delprim and *Melinis minutiflora* plants grown in a greenhouse. The absolute peak areas were divided by the area of the internal standard peak and z-score was calculated (peak area - mean peak area/standard deviation of peak). The dendrogram of compounds was constructed via hierarchical clustering based on Euclidean distances. The major volatile constituents of intact *D. intortum* headspace were 2-heptanone and 3-heptanone. Monoterpenoids were only detectable after 48 hrs of *S. frugiperda* feeding, when (*E*)-4,8-dimethyl-nona-1,3,7-triene ((*E*)-DMNT), (*Z*)-*β*-ocimene, (*E*)-*β*-ocimene and (*E*)-alloocimene were emitted. The relative (*E*)-DMNT emission, (*E*)-*β*-ocimene emission and total monoterpenoid emission of intact and herbivore induced *D. intortum* were significantly different in pairwise comparisons with Kruskal-Wallis tests and pairwise comparisons with Wilcoxon rank sum test with Benjamini and Hochberg p-correction (*χ^2^* = 57.315, p = 0.00012, *χ^2^* = 52.321, *p* = 8.5*10^-5^, and *χ^2^* = 52.904, *p* = 7.74*10^-4^). Linalool, *β*-myrcene were present in the headspace of intact maize. In response to 48 hrs of larval feeding (*E*)-DMNT, (*Z*)-*α*-bergamotene, *β*-caryophyllene, (*Z*)-*β*-farnesene, humulene and *β*-bisabolene were emitted. The relative (*E*)-DMNT emission and total sesquiterpenoid emission of intact and herbivore induced *Z. mays* cv. Delprim was significantly different using the same statistical tests (*χ*^2^ = 57.315, *p* = 3.1*10^-4^ and *χ^2^* = 59.163, *p* = 8.2*10^-4^). The volatile headspace of the both intact and herbivore-induced *M. minutiflora* is composed of a variety of monoterpenoid and sesquiterpenoid compounds, such as (*E*)-DMNT, limonene, germacrene-D. Neither the relative (*E*)-DMNT emission nor the total monoterpenoid emission nor the total sesquiterpenoid emission of intact and herbivore induced *M. minutiflora* were significantly different in the same statistical tests (*χ^2^* = 57.315, *p* = 0.62, *χ*^2^ = 52.904, *p* = 0.63 and *χ*^2^ = 59.163, *p* = 0.12).

**Fig. 3:**
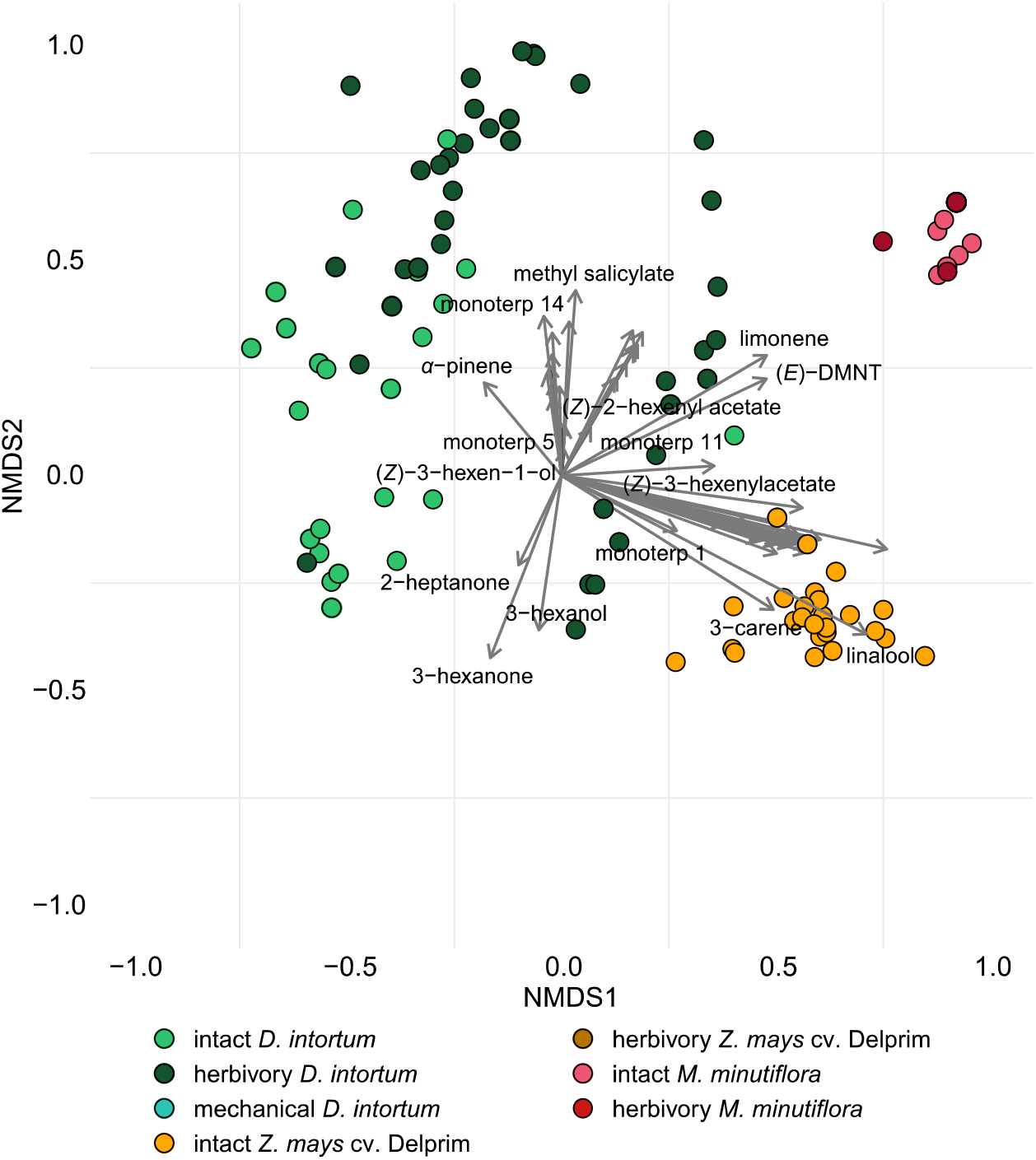
Ordination of volatile samples from intact, herbivore damaged and mechanically damaged *Desmodium intortum, Zea mays* cv. Delprim and *Melinis minutiflora* plants based on non-metric multidimensional scaling (NMDS). The NMDS plots were based on presence-absence values and calculation of Jaccard-dissimilarity indices. The stress value of the plot is 0.138. Vectors represent correlations of volatile features with distribution of plant samples along the NMDS1 and NMDS2 axes.

**Fig. 4:**
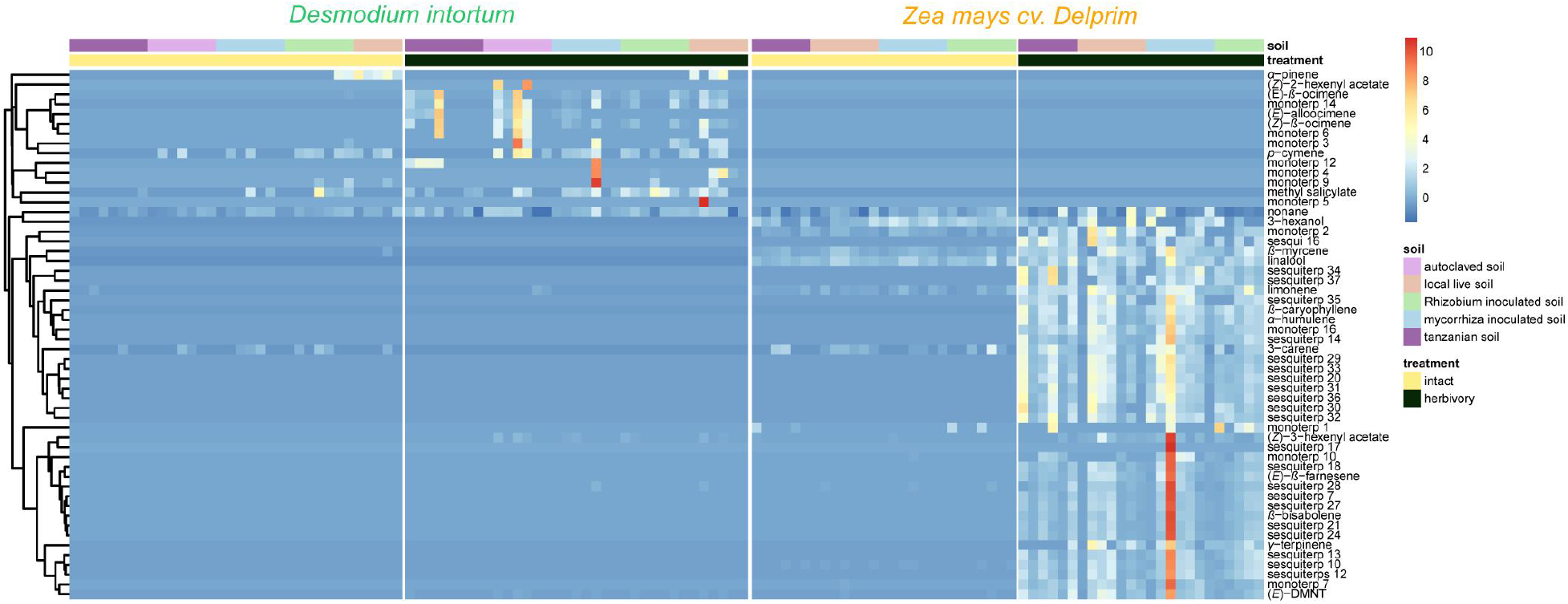
Volatile emission profile of intact and herbivore damaged *Desmodium intortum* and *Zea mays* grown in soils with different microbial composition. The absolute peak areas were divided by the area of the internal standard peak and z-score was calculated (peak area - mean peak area/standard deviation of peak). The dendrogram of compounds was constructed via hierarchical clustering based on Euclidean distances.

**Fig. 5:**
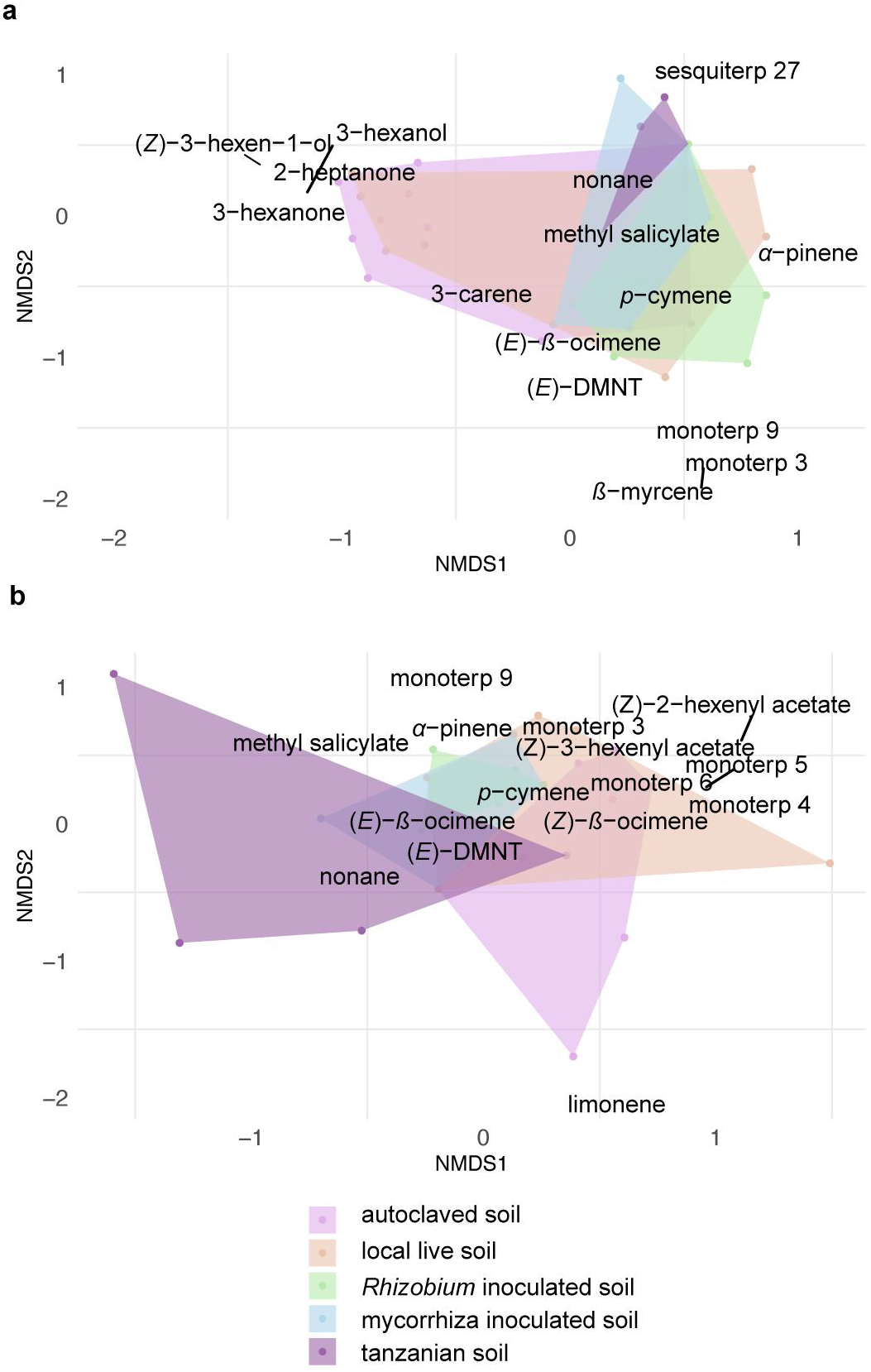
The absence of volatile terpenoids in intact *Desmodium intortum* does not result from poor soil microbiota and insufficient nodulation. **a,** Non-metric multidimensional scaling (NMDS) ordination of volatile profiles from headspace of intact plants. **b,** NMDS ordination of herbivore-damaged *D. intortum* plants grown in different soils in a greenhouse. The stress values of NMDS ordination were 0.146 for intact and 0.120 for herbivore induced plants. The volatile profile of intact *D. intortum* on different soil treatments largely overlap while upon herbivory, some differentiation is observed. Scaling is based on Jaccard-distance matrix calculated from centered area values for each compound. The stress values are 0.146 and 0.120 for NMDS ordination of intact and herbivore-induced samples. Based on PERMANOVA and pairwise comparison of plants grown in different soil treatments the volatile profile of intact (F_model_ = 3.260, R^2^ = 0.189,*p*_adj_ = 0.615) and herbivore-induced *D. intortum* (F_model_ = 7.268, R^2^ = 0.326, *p*_adj_ = 0.090) did not cluster separately.

**Fig. 6:**
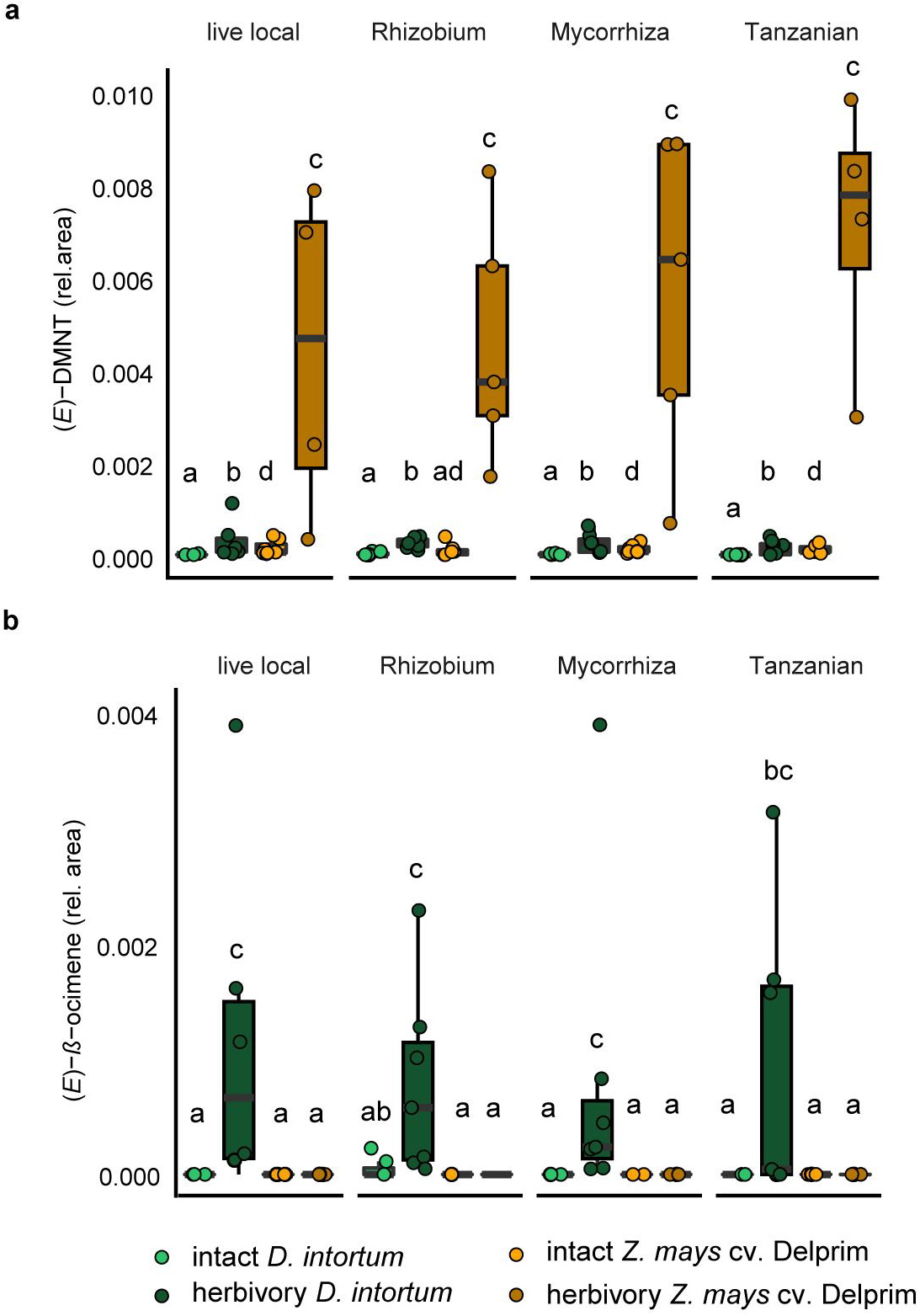
The emission profile of *Desmodium intortum* and *Zea mays* cv. Delprim was not significantly altered by soil microbial treatments. **a,** The relative (*E*)-4,8-dimethyl-nona-1,3,7-triene ((*E*)-DMNT) emission and (*E*)-*β*-ocimene emission of *D. intortum* and *Z. mays* cv. Delprim plants grown in soils containing *Rhizobium* spp., mixture of mycorrhizal fungi and soil of push-pull fields. The absolute peak areas were divided by the area of the internal standard peak to calculate relative values. The error bars show the standard error in relative emission units. Inoculation did not alter significantly the relative (*E*)-DMNT (*χ^2^* = 80.156, *p* = 0.303). **b,** Neither did inoculation affect the (*E*)-ß-ocimene (*χ*^2^ = 7.688, *p* = 0.103) emissions of intact *D. intortum* plants based on pairwise comparisons with Kruskal-Wallis test with Wilcoxon rank sum test with Benjamini and Hochberg p-correction. Herbivore induced *D. intortum* plants grown in different soils were also not significantly different from each other in the relative (*E*)-DMNT (*χ^2^* = 5.153, *p* = 0.272) and (*E*)-*ß*-ocimene (*χ^2^* = 80.395, *p* = 0.268) emissions.

**Fig. 7:**
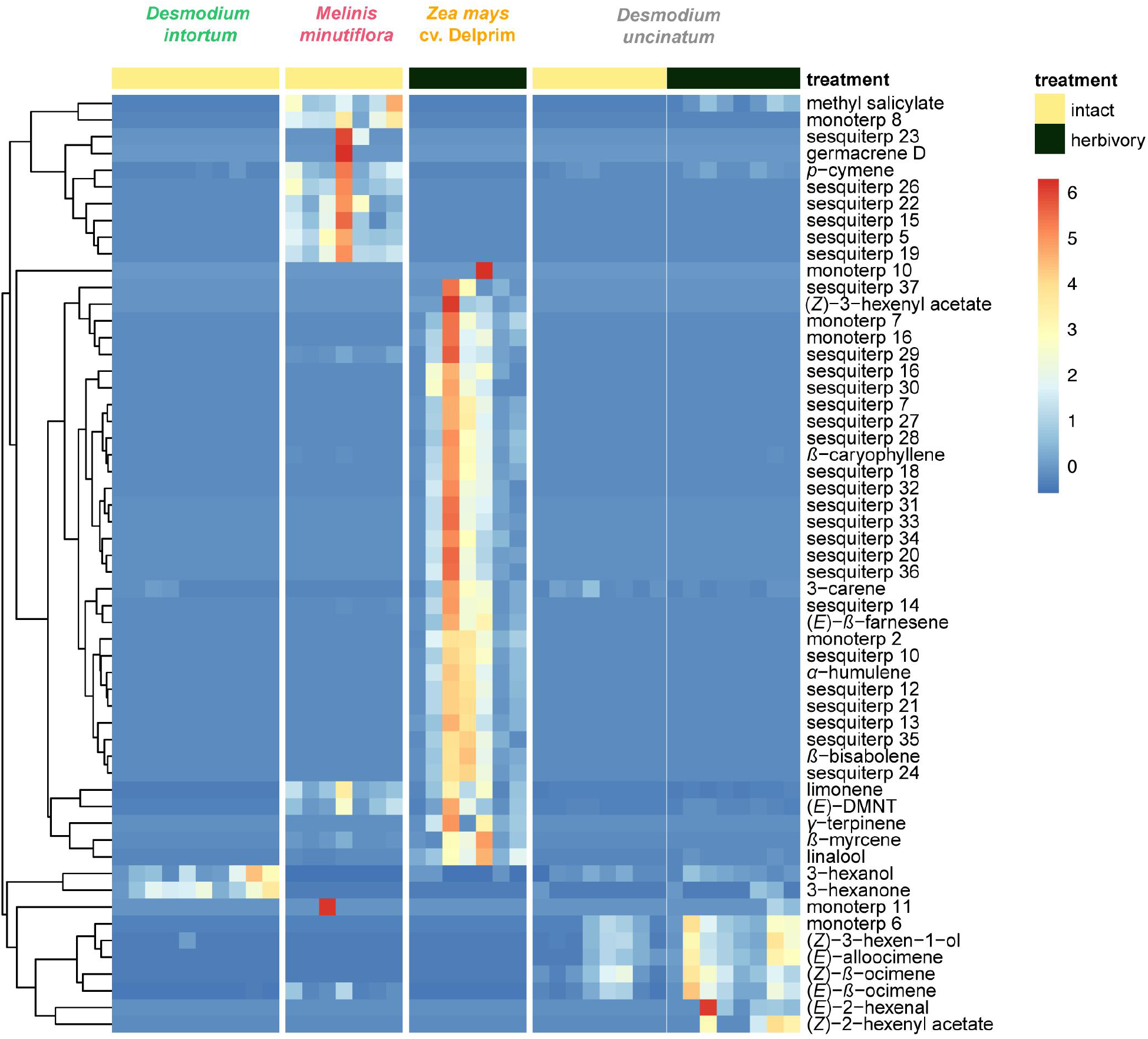
Volatile emission of *Desmodium uncinatum* and *Desmodium intortum* compared to *Melinis minutiflora* and *Zea mays* cv. Delprim. The heatmap shows the relative amounts of volatile compounds emitted from intact *D. intortum* (greenleaf *Desmodium*), *M. minutiflora* and *D. uncinatum* (silverleaf *Desmodium*) as well as herbivore-damaged *Z. mays* (maize) and *D. uncinatum* plants. The absolute peak areas were divided by the area of the internal standard peak and z-score was calculated (peak area - mean peak area/standard deviation of peak). The dendrogram of compounds was constructed via hierarchical clustering based on Euclidean distances.

**Fig. 8:**
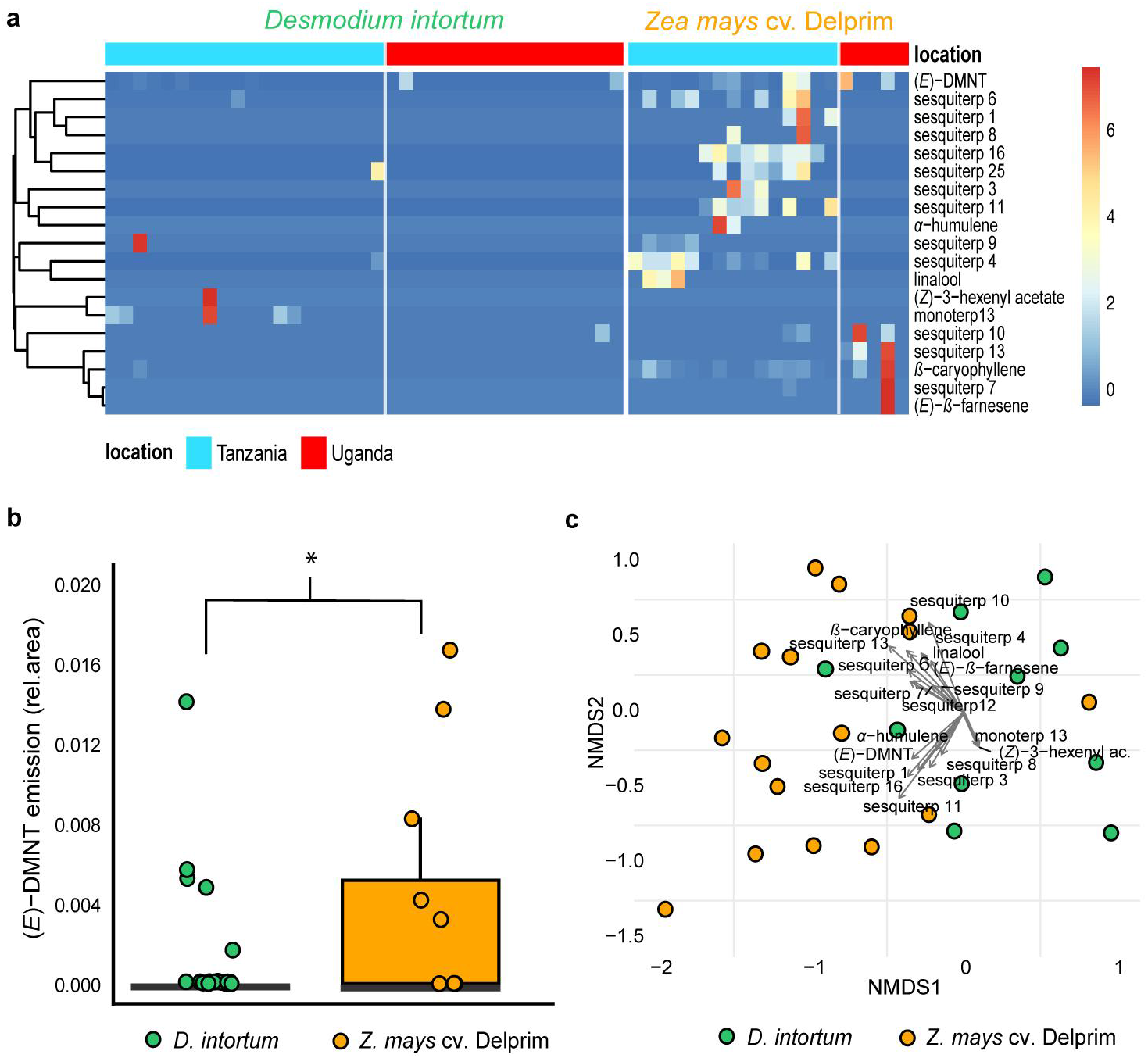
Volatile emission of field grown *Desmodium intortum* and *Zea mays* plants from two locations. **a,** Heatmap volatile emissions of *D. intortum* (greenleaf *Desmodium*) and *Z. mays* plants at locations in Tanzania and Uganda. The absolute peak areas were divided by the total area of compounds belonging to monoterpenoids, sesquiterpenoids or green leaf volatiles per location and z-score was calculated (peak area - mean peak area/standard deviation of peak). The dendrogram of compounds was constructed via hierarchical clustering based on Euclidean distances. **b,** Similarly to greenhouse experiment, the constitutive emission of monoterpenoids, such as (*E*)-4,8-dimethyl-nona-1,3,7-triene ((*E*)-DMNT) and (*E*)-*β*-ocimene were not detectable in case of *D. intortum* plants, due to possible underlying biotic and abiotic stressors emission of (*E*)-DMNT was visible in a small fraction of *D. intortum* samples. Based on Kruskal-Wallis tests and Wilcoxon rank sum test with Benjamini and Hochberg p-correction the relative (*E*)-DMNT abundance of *Z. mays* volatile samples was significantly higher than that of *D. intortum* volatile samples (*χ^2^* = 15.310, *p* = 2*10^-3^). **c,** Non-metric multidimensional scaling (NMDS) of the volatile profile of *D. intortum* and *Z. mays* plants from field locations. The vectors represent the correlation of volatile features with the distribution of plant samples along the NMDS1 and NMDS2 axes. The stress value of the NMDS plot is 0.116. Based on PERMANOVA and pairwise comparison the volatile profile of *D. intortum* and *Z. mays* were significantly different (F_model_ = 8.816, R^2^ = 0.149, *p*_adj_ = 1*10^-3^).

**Fig. 9:**
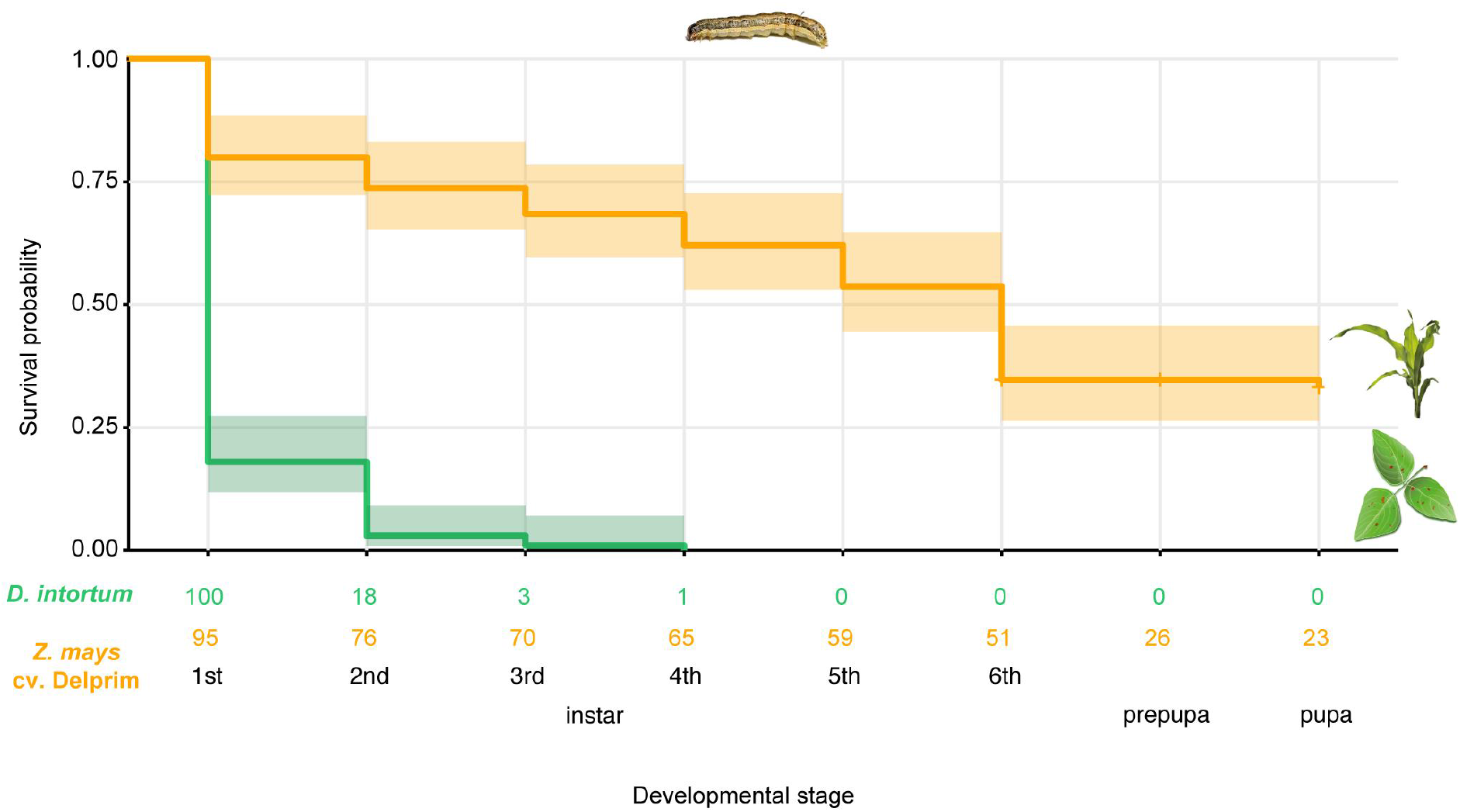
The survival probability of *Spodoptera frugiperda* on diets consisting of *Desmodium intortum* (greenleaf *Desmodium*) or *Zea mays* cv. Delprim (maize) leaves. The Kaplan-Meier survival curves show that larvae on *D. intortum* diet had significantly higher mortality than larvae on *Z. mays* diet (*p* = 2*10^-16^). The *D. intortum* diet resulted in a total mortality by the 4th instar larval stage. The inset below the plot shows the number of specimens reaching each developmental stage on the two types of diets.

**Fig. 10:**
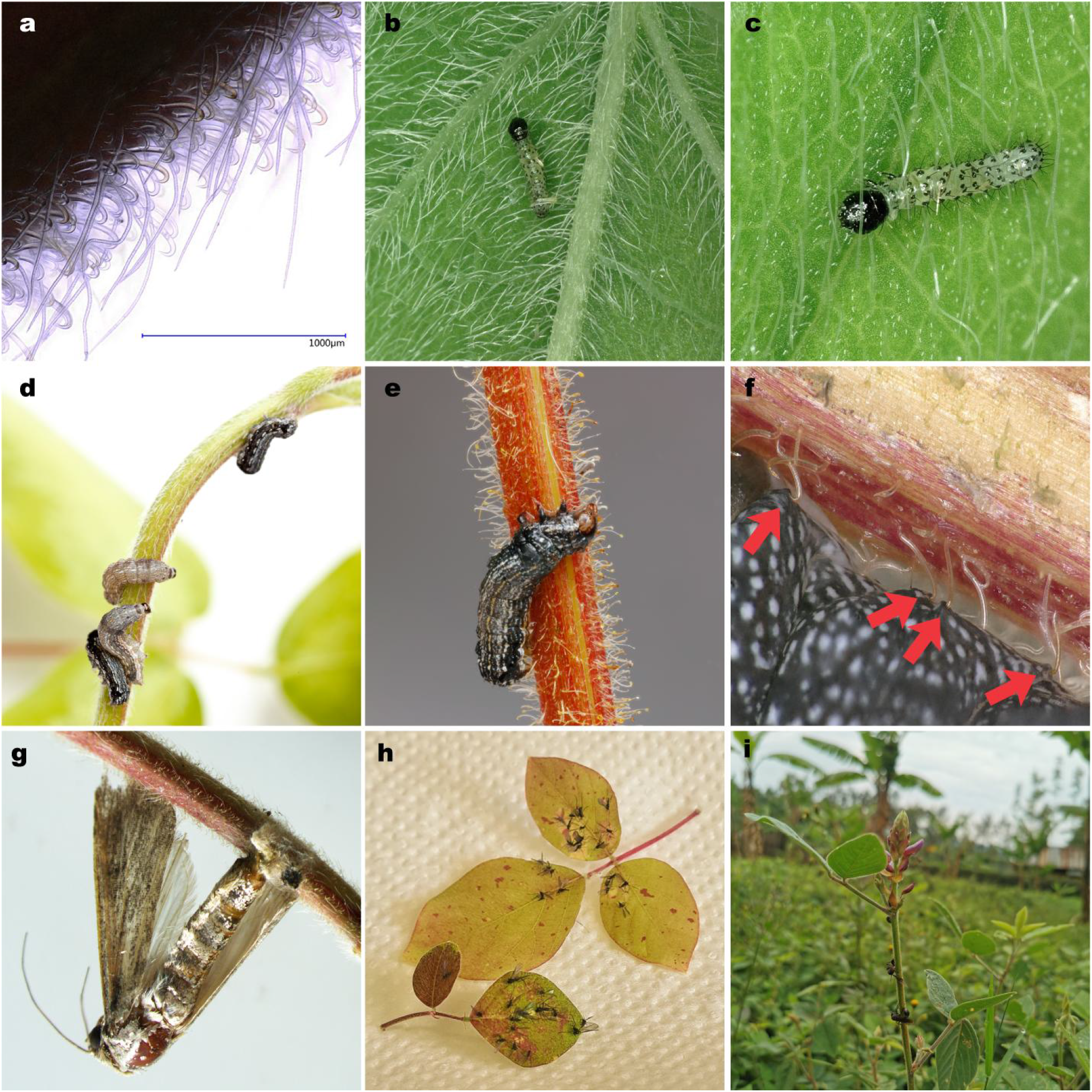
*Spodoptera littoralis* larvae and adult *Spodoptera frugiperda* immobilized on *Desmodium intortum* and *Desmodium uncinatum* stems. **a,** Light microscopic picture of trichomes on the stem of *D. intortum*. **b-c,** Despite the dense network of sharp, straight and hooked trichomes, neonate larvae of *Spodoptera* spp. are able to graze and easily navigate through the leaf surfaces of *D. intortum*. **d-e**, Immobilized *S. littoralis* larvae on stems of *D. uncinatum* and on *D. intortum* stems. **f,** The cuticle of an *S. littoralis* larva pierced by uncinate trichomes, the red arrows indicate puncture sites. **g,** Ovipositing *S. frugiperda* female immobilized on *D. intortum*. **h,***Bradysia sp*. immobilized on *D. intortum* leaves. **i,** *Hymenopteran* insects immobilized on *D. intortum* stems at a volatile collection site in Mwanza, Tanzania.

## SUPPLEMENTARY INFORMATION

### Chemical analysis of GC-MS samples

Compounds were tentatively identified by matching their mass spectra with those found in MS Libraries (NIST11 and Wiley). The identification was verified by synthetic standards and matching Kovats retention indices found in literature for DB-WAX and HP-5 capillary columns.

Retention indices of volatile components, GC-MS raw data, the list of synthetic standards (suppliers and purity information) that were injected on DB-WAX and HP-5 columns to verify library based identification of headspace volatile components and behavioral bioassay data are available on FigShare (https://figshare.com/account/projects/134051/articles/19297730).

### Volatile collections site selection

Samples were collected from Tarime and Musoma districts in Mara region in Tanzania. A small survey of the farmers practicing push-pull farming and/or growing *Desmodium* spp. was conducted to identify suitable sampling sites. Four locations were selected for *Desmodium intortum* volatile collection, three of them being *D. intortum* (greenleaf desmodium) monoculture and one of them a push-pull plot. One farm was selected for collection of volatiles also from herbivore infested maize plants.

*Desmodium* intortum volatiles samples were also collected in Uganda during the rainy season from Rural Community in Development (RUCID) centre, in Mityana district, from a *D. intortum* monoculture. The plots had been growing for four years at the time of sampling and were managed by trimming *Desmodium* about six times a year to feed animals.

| Sampling site | Practice | GPS coordinates | Relative humidity | Temperature |
| --- | --- | --- | --- | --- |
| Kitagasembe village, Gwitiriyo ward, Tarime district, Mara region, Tanzania | *D. intortum* monoculture, Maize with common beans as intercrop (*Phaseolus vulgaris*) | -1.3, 34.4792 | 70% | 20 - 22° C |
| Gwitiriyo, Gwitiriyo ward, Tarime district, Mara region | *D. intortum* monoculture | -1.266661, 34.488133 | 68% | 20 - 22° C |
| Kyoruba village, Pemba Ward, Tarime District. | *D. intortum* monoculture | -1.318, 34.520 | 75% | 20 - 22° C |
| Vi Agroforestry center, Lubango Ward, Musoma District, Mara region | Push-pull farming with *D. intortum* intercropping | -1.53054, 33.857955 | 81% | 20-22° C |
| RUCID centre, Mityana district, Uganda | *D. intortum* monoculture and maize monoculture | 0.437941, 32.042500 | 68% | 20- 26° C |

### Visualization of wind tunnel oviposition bioassay

The three dimensional model of wind tunnel assays (Extended Data Fig.1) was prepared in SketchUp (version 20.0).

